# Learning Feedback Molecular Network Models Using Integer Linear Programming

**DOI:** 10.1101/2021.11.08.467837

**Authors:** Mustafa Ozen, Ali Abdi, Effat S. Emamian

## Abstract

Analysis of intracellular molecular networks has many applications in understanding of the molecular bases of some complex diseases and finding the effective therapeutic targets for drug development. To perform such analyses, the molecular networks need to be converted into computational models. In general, network models constructed using literature and pathway databases may not accurately predict and reproduce experimental network data. This can be due to the incompleteness of literature on molecular pathways, the resources used to construct the networks, or some conflicting information in the resources. In this paper, we propose a network learning approach via an integer linear programming formulation that can efficiently incorporate biological dynamics and regulatory mechanisms of molecular networks in the learning process. Moreover, we present a method to properly take into account the feedback paths, while learning the network from data. Examples are also provided to show how one can apply the proposed learning approach to a network of interest. Overall, the proposed methods are useful for reducing the gap between the curated networks and experimental network data, and result in calibrated networks that are more reliable for making biologically meaningful predictions.

## 1 Introduction

Molecular networks are the networks of biochemical interactions between the molecules. They can be portrayed as directed graphs in which nodes represent biological molecules, i.e., proteins, genes etc., and edges represent biochemical interactions between the molecules [1–4]. Research and development of such networks has application in target discovery and drugs development, and analyzing the role of the molecular component in disease pathogeneses [5, 6], understanding cellular decision making processes [7, 8], understanding cell development and cell differentiation [9], developing molecular fault diagnosis and signaling capacity analysis methods [10–13], identifying disease subtypes and their regulators [14], and many other applications for better understanding of human diseases. Hence, constructing and analyzing molecular network models have emerged particularly over the past decade as an important area of systems biology research.

To study molecular networks, one needs to convert the molecular network graphs into computational models so that they can be analyzed to obtain novel and biologically relevant results. Continuous and discrete models have been widely studied so far. One way to model molecular networks is to convert them into a mathematical form, by building a system of differential equations that can capture temporal and spatial behaviors of molecules within a complex network. A main challenge in this approach is that the mechanistic details and kinetic parameters of the molecular interactions are not available for continuous models in large molecular networks. In such scenarios, Boolean modeling has been useful as it does not need detailed kinetic information and still can provide biologically relevant results, as discussed in [10–13], and in several review articles [15–24] that are summarizing many other original research contributions.

Typically, models for literature-curated molecular networks do not adequately match experimental data. This is due to the incompleteness and species heterogeneity of resources, databases, and the literature used to construct the networks. In such networks, for some individual interactions, generally there is more than one publication, and sometimes some studies suggest contradicting results. Consequently, models constructed for molecular networks using only the literature may poorly perform in terms of fitting experimental data. Thus, one needs to develop algorithms to learn and refine the models, so that the learned networks can mimic the experimental observations [5, 25–27]. Herein, we propose a method for fitting a network model to data. The method transforms the model into an integer linear programming (ILP) formulation, allowing us to learn a subnetwork of the initial network that exhibits an optimal fit to the experimental data. As discussed in what follows, the method incorporates the role of network regulatory feedback mechanisms, not considered in prior studies [5, 26, 27].

Modeling and analysis of molecular networks become more challenging if there are positive or negative feedback paths in the network. Due to the feedback mechanisms, network responses may change over time because of some compensatory or regulatory mechanisms [28, 29]. Feedbacks can cause delays in propagation of signals to the network outputs, while passing through the feedback paths. Therefore, incorporating the delays caused by the feedbacks, which may result in different network responses over time, is essential when developing network learning algorithms. The goal of this paper is to introduce new network learning ILP formulations for different Boolean models, when the network of interest has some feedback paths, in addition to the feedforward paths.

The rest of this paper is organized as follows: In Section 2, we present two Boolean models, along with their truth tables and examples. In Section 3, we present an ILP formulation for each model, then demonstrate in Section 4 using some numerical examples, how networks with feedbacks can be learned from data. Finally, we conclude the paper with some remarks given in Section 5.

## 2 Molecular Network Models

Fig. 1 illustrates a toy molecular network having 7 nodes and 10 edges. Each edge represents an interaction between the molecules. An arrow edge “→” represents an *activatory* interaction and a blunt edge “—|” reflects an *inhibitory* interaction. A node at the beginning of an edge represents an input molecule, whereas a node at the end of an edge stands for a product (output). A set of input nodes and a product node together constitute an *interaction set*. For instance, in Fig. 1, the nodes B, G, C and E together represent an interaction set in which B, G and C are the input molecules and E is the output molecule (product). Overall, it can be said that molecular networks consist of interaction sets, each set comprising one or more inputs and one output. A molecule is defined as *active* if its abundance or activity level, e.g., phosphorylation level, is above a biologically significant threshold, and *inactive* otherwise.

**Fig. 1.**
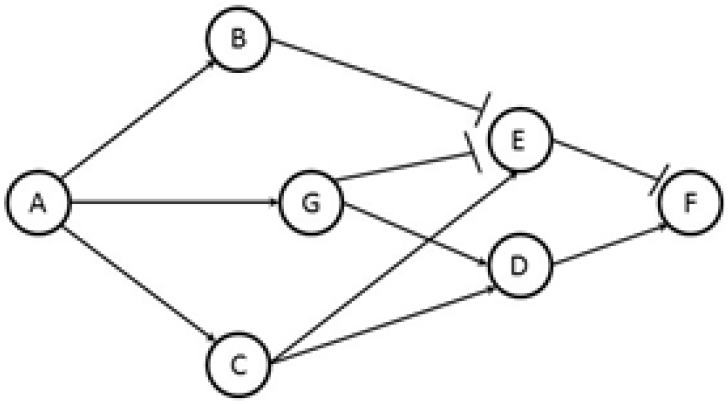
Toy example of a molecular network.

To model how the output molecule of each interaction set is controlled by its input molecules, 0 and 1 values and logic operations are used in the Boolean framework [10, 13, 15, 16]. A key advantage of this framework is that it does not need the detailed mechanistic information about the molecular interactions in various interaction sets, i.e., it does not have hundreds of unknown parameters to be estimated, and yet provides certain biologically meaningful results and predictions. In the rest of this section, two such models, Model I and Model II, are presented, and then are used in subsequent sections, for the network modeling and learning.

### 2.1 Model I: 1 for Increased Activity and 0 for Decreased or Not Changed Activity

In a typical biological interaction set, the activity level of the output, the product of the interaction set, can increase, decrease, or remain unchanged, compared to its basal level and depending on its input molecules. In Model I, *increase* in the activity level of a molecule is represented by a 1, whereas *decrease* or *no change* in the activity level of a molecule is represented by a 0. Assume there exists an interaction set with multiple activators and inhibitors. Model I incorporates the following two rules to specify the output molecule’s activity level: Let *x*_1_,…, *x*_*i*_, *x*_*i*+1_,…, *x*_*n*_ be *n* input molecules and *w* be the product of an interaction set such that *x_i_* is an activator for *i* = 1,…, *j* and *x*_*i*_ is an inhibitor for *i* = *j* + 1,…, *n*. Then,

a. If at least one of the activators and none of the inhibitors are 1, i.e., active, then the output is 1. This means if ⴺ *i* ∈ {1,…, *j*} such that *x_i_* = 1 and *x_i_* = 0 for all *i* = *j* + 1,…, *n*, then *w* = 1.
b. If at least one of the inhibitors is 1, then the output is 0, i.e., if ⴺ *i* ∈ *j* + 1,…, *n* such that *x_i_* = 1, then *w* = 0.

Fig. 2 is an illustration of the model and its truth table, based on its rules (a) and (b) given above.

**Fig. 2.**
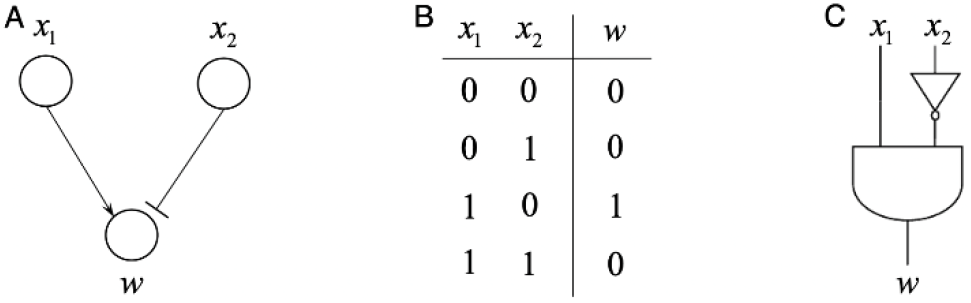
An example for Model I. (A) A two-input one-output interaction set. (B) Truth table of the interaction set based on the model rules. (C) Logic circuit representation of the interaction set using NOT and AND gates.

### 2.2 Model II: 1 for Changed Activity and 0 for Not Changed Activity

In Model II, *change* and *no change* in the activity are considered, and are labeled as 1 and 0, respectively. In response to an input signal, we declare a *change* if the activity of a molecule increases or decreases, compared to its basal level. On the other hand, we declare a *no change* if the activity of a molecule does not change with respect to its basal level, when an input signal is applied.

Recall that *x*_1_,…, *x_i_*, *x*_*i*+1_,…, *x_n_* are the *n* input molecules and *w* is the product of an interaction set, such that *x_i_* is an activator for *i* = 1,…, *j* and *x_i_* is an inhibitor for *i* = *j* + 1,…, *n*. In Model II, each interaction set in the network follows these two rules:

a. The output is 1, meaning that there is a change in the output’s activity, if at least one of the input molecules is 1, i.e., if ⴺ *i* ∈ {1,…, *n*} such that *x_i_* = 1, then *w* = 1.
b. The output is 0, meaning that there is no change in the output’s activity, if all the inputs are 0, i.e., *w* = 0 if *x_i_* = 0 for all *i* = 1,…, *n*.

The model and its truth table are exemplified in Fig. 3, using its two rules given above. Overall, this model incorporates and reflects any changes in the input of an interaction set as a change at its product.

**Fig. 3.**
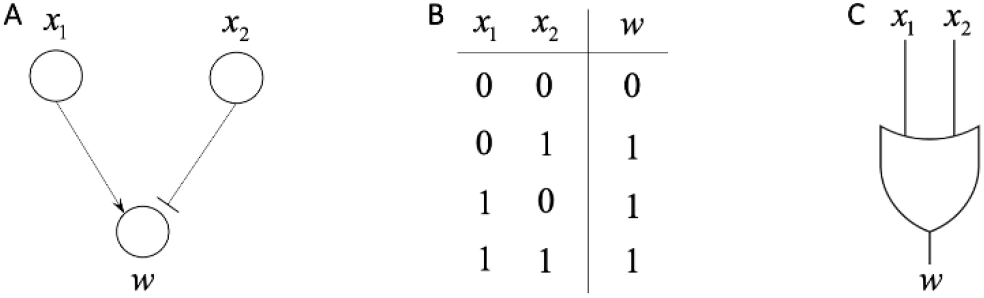
An example for Model II. (A) A two-input one-output interaction set. (B) Truth table of the interaction set based on the model rules. (C) Logic circuit representation of the interaction set using an OR gate.

## 3 Learning Molecular Network Models

As discussed in Section 1, model predictions of literature-based networks may not agree with experimental data. In order to fit the network to the experimental data, some of the edges (interactions) in the network may need to be removed (spurious interactions), or some new edges may need to be added (missed interactions), so that the resulting network can reflect actual collective behaviors of the molecules, i.e., models that fit the experimental data. Herein, we focus on removing edges and finding a sub-network of the initial network, since adding new edges requires having access to additional original publications or performing many experiments that are costly and time consuming to acquire. One way to do this is to conduct an optimization to remove edges one by one and check the number of mismatches between model predictions and experimental data. However, for large networks, this does not help as removing one edge at a time most often does not change model predictions. For this reason, we convert this problem into an ILP problem so that multiple edges can be removed systematically, and finally a subnetwork of the initial network can be found as the optimal solution that fits the data. A similar approach is studied in [5] on a network that does not include feedbacks. In this paper, we present a method in Section 4 that incorporates feedback interactions in network learning, after presenting our ILP formulations for Models I and II in this section.

Our goal is to minimize the number of mismatches between model responses and the experimental data. The experimental data set is typically obtained by treating cells with selective agonist of the input molecules and then measuring the activity, i.e., the protein or phosphoprotein levels, of some of the intermediate and output molecules by running western blot analysis.

Let *n_E_* be the number of experiments and each experiment be indexed by the superscript *k* = 1,… *n_E_*. In the network, assume there exist *n_R_* interaction sets that are indexed by the subscript *i* = 1,…, *n_R_*. Each interaction set *i* has the corresponding index set *I_i_* = *A_i_* ∪ *H_i_* for its input molecules, in which *A_i_* and *H_i_* are the index sets of activators and inhibitors, respectively. Lastly, let *M* be the index set of molecules for which we have experimental data. Then, in the general form of the proposed ILP formulation, we define all the other variables as shown below.

- 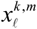 : experimental value of the *ℓ*^th^ node in the *k*^th^ experiment, for all indicates that the *ℓ* ∈ *M*. Here, the superscript *m* indicates that the *ℓ*^th^ node has experimental measurement in the *k*^th^ experiment.

Then, in each interaction set *i*, we have:

- 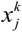 : model’s predicted value of the *j*^th^ input node of the *i*^th^ interaction set in the *k*^th^ experiment, for all *j* ∈ *I_i_*. To simplify the notation, the interaction set label *i* is not included in the 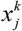 variable.
- *y_j_* : decision variable, for all *j* ∈ *I_i_*. *y_j_* = 1 means that *j*^th^ edge in the interaction set *i* should be preserved in the network whereas *y_j_* = 0 means that *j*^th^ edge in the interaction set *i* should be removed from the network.
- 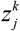 : transition variable, for all *j* ∈ *I_i_*. It transits the input value 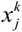 associated with the *j*^th^ edge to the output of interaction set *i*, i.e., 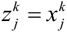, if *y_j_* = 1. Otherwise, 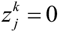.
- 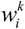 : output (product) of the interaction set *i* in the experiment *k*.

The objective function to be minimized in the learning phase is the summation of the mismatches - absolute differences - between the experimental data and model’s predictions over all experiments. Thereby, the objective function is:

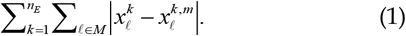

For binary 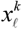 and 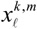 values, (1) can be linearized as:

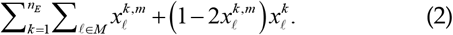

### 3.1 ILP Formulation for Model I

Using all the definitions given above, the constrained ILP formulation for Model I introduced in Section 2.1 can be written as follows:

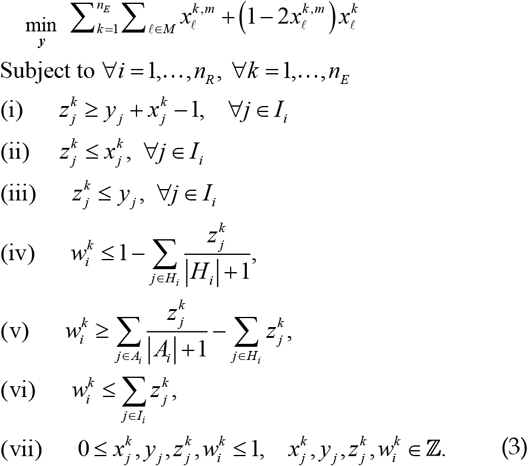

In (3), ***y*** = [*y_j_*] is the vector of indices of edges in the network, and the constraints (i), (ii), and (iii) are introduced for edge removal. More precisely, these three constraints assure that if the *j*^th^ interaction in interaction set *i* is removed, i.e., *y_j_* = 0, then the transition variable is 0, i.e., 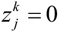, so that the input molecule associated with the *j*^th^ interaction does not affect the value of the interaction set output 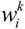. If the *j*^th^ interaction needs to stay, i.e., *y_j_* = 1, then these constraints guarantee for the transition variable 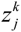 that 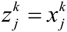. The constraints (iv), (v), and (vi) implement the two rules of Model I. To elaborate, depending on the constraints (i), (ii), and (iii), the transition variable 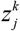 becomes either 0 or 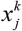. Then, if none of the inhibitors and at least one of the activators is 1, the constraints (iv) and (v) guarantee that the interaction set output 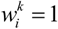 (Section 2.1, rule (a)). On the other hand, if at least one of the inhibitors is 1, i.e., ⴺ*j* ∈ *H_i_* such that 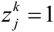, then, the constraints (iv) and (v) make sure that the interaction set output 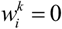 (Section 2.1, rule (b)). The constraint (vi) is necessary to guarantee that the interaction set output 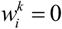, if all the incoming edges are removed or the input values of the remaining edges are 0. Lastly, the constraint (vii) is needed to guarantee that all variables are integers, and they are either 0 or 1.

### 3.2 ILP Formulation for Model II

A similar formulation can be developed for learning Model II introduced in Section 2.2. This can be done by discarding the constraint (iv) of (3) and replacing the constraint (v) of (3) by 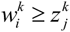 for all *j* ∈ *I_i_*, which result in the following constrained ILP formulation for Model II:

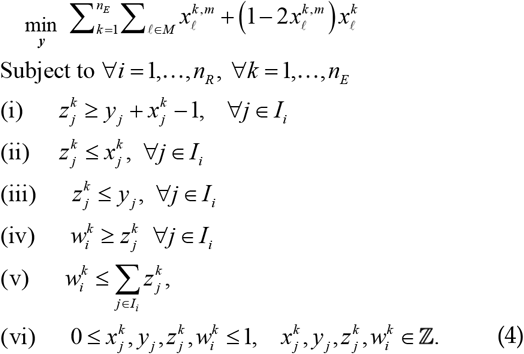

Similarly to (3), ***y*** = [*y_i_*] in (4) is the vector of indices of edges in the network, and the constraints (i), (ii), and (iii) in (4) are introduced for edge removal, as elaborated in the previous subsection. The constraints (iv) and (v) in (4) implement the two rules of Model II. More precisely, if at least one of the input values whose associated edge is not removed is 1, then we have 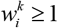 from (iv) and (v), which guarantees 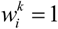, because of the constraint (vi). Otherwise, 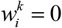. Finally, the constraint (vi) is needed to guarantee that all variables are integers, and they are either 0 or 1.

## 4 Numerical Results

The ILP formulations in (3) and (4) search for a vector ***y*** = [*y_j_*], the vector of indices of edges in the network, to minimize the number of mismatches between predictions and the data. To be more precise, a network can be represented by the vector *y* that is a vector of 1s whose length is equal to the total number of interactions in the network. Thus, a subnetwork of the initial network can be represented by the same length *y* vector where some of the 1s there are changed to 0 (if the *j*^th^ entry of *y* is 0, then this means that the *j*^th^ interaction is not present in the sub-network). As a result, by solving the ILP formulations, one can find the best *y* vectors, i.e., the subnetworks, that have the optimal fit to the data.

### 4.1 An Exemplary Network

Now we apply the ILP formulation in (3) to the exemplary network in Fig. 4A. The equations - based on the two rules of Model I - for each node can be written as shown in Fig. 4B for Model II, the ILP formulation in (4) has to be used). Because of the presence of feedback interactions, the blue edges in Fig. 4A, this network has two early event (EE) and late event (LE) representations, as shown in Fig. 5A and 5D. This is because when there are feedbacks in the network, the network response can be different at different time instances, due to the signal propagation delays caused by the feedback paths. Using the equations in Fig. 5B and 5E for EE and LE networks, respectively, EE and LE truth tables can be created, as given in Fig. 5C and 5F.

**Fig. 4.**
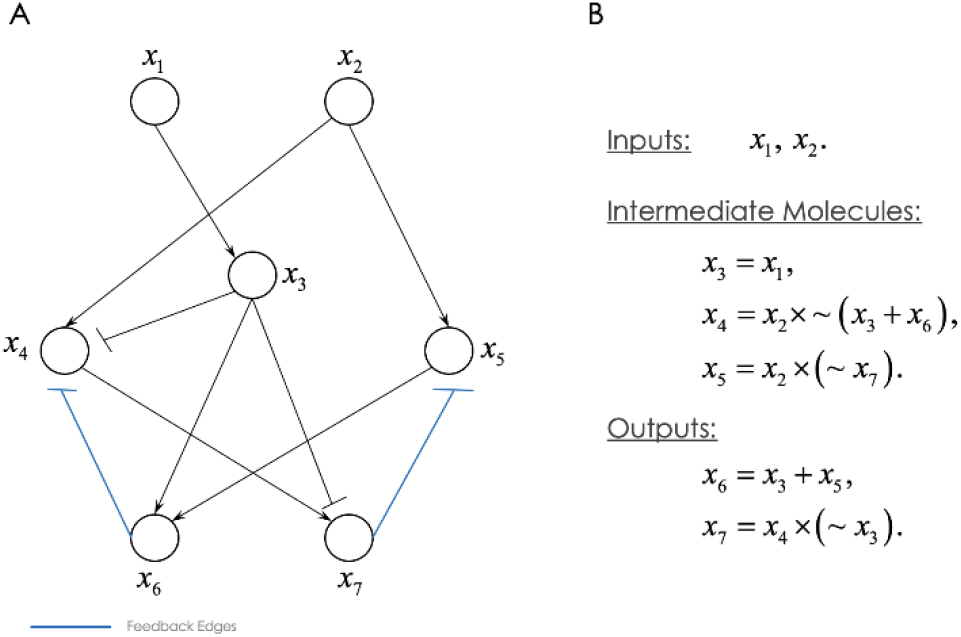
An exemplary network and its equations. (A) The network with two input nodes and two output nodes, where feedforward and feedback edges are shown in black and blue, respectively. (B) The equations for the nodes obtained using Model I.

**Fig. 5.**
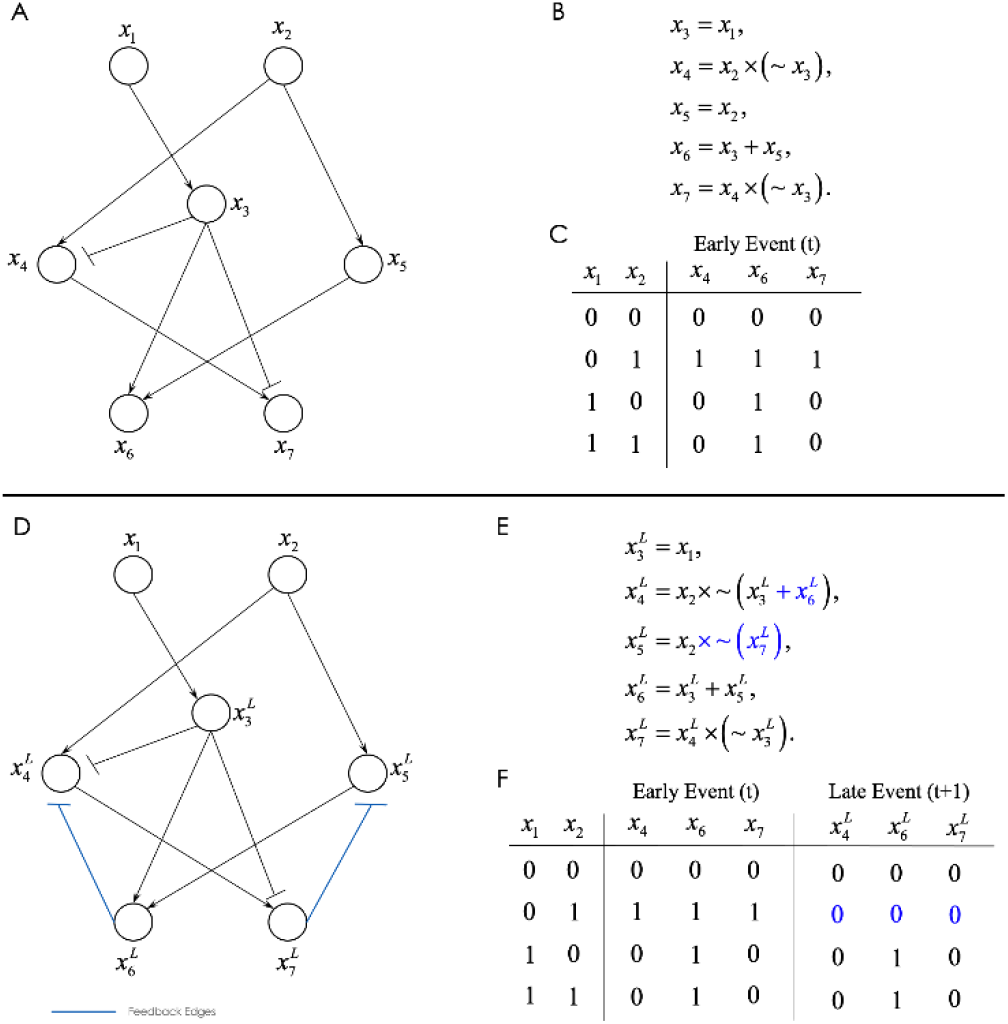
The early event (EE) and the late event (LE) representations of the exemplary network. (A) The EE network. (B) Equations of the nodes in the EE network. (C) Truth table of the EE network. (D) The LE network. (E) Equations of the nodes in the LE network. (F) Truth table of the LE network (truth table of the EE network is also included for convenience).

### 4.2 Network Learning from Data

Assume that the network in Fig. 5D is the ground truth network and Fig. 5F is its truth table that is obtained via hypothetical lab experiments, i.e., measuring early and late output activities in response to different input activities. To test the ILP formulation in (3), we alter this network by adding some spurious edges and generate a new network shown in Fig. 6A, that has a new truth table given in Fig. 6B.

**Fig. 6.**
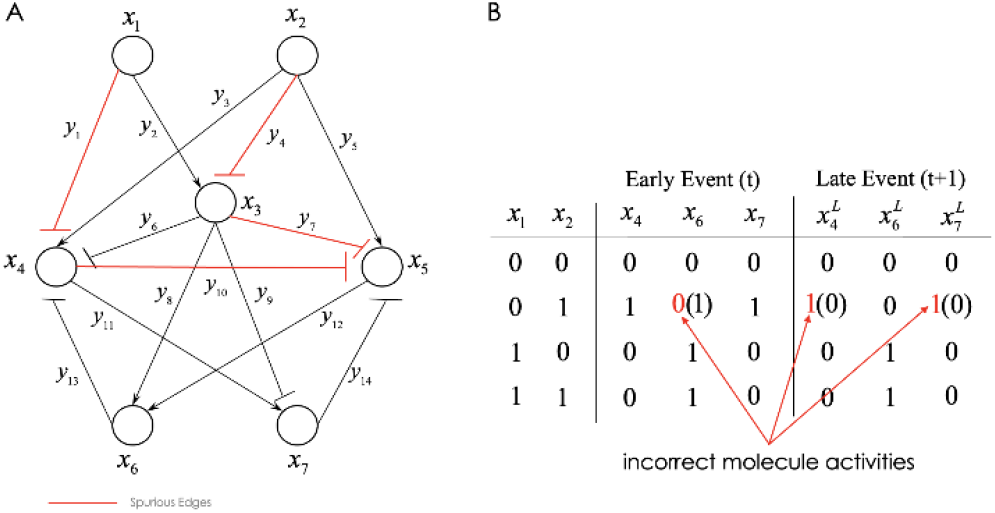
The exemplary network with some spurious edges and its incorrect truth table. (A) Red edges in the network represent spurious interactions that are – unknowingly – included during the network curation from the literature, in addition to the correct interactions shown by black edges. (B) Red entries in the truth table represent incorrect activity levels, caused by the spurious interactions, whereas the corresponding correct activity levels are given in parentheses.

Suppose that the new network with the spurious interactions (Fig. 6A) is the initial network constructed from literature, and therefore our initial truth table has some mismatches, the red values in Fig. 6B, compared to the experimental truth table in Fig. 5F. Our goal is to learn this network from the experimental data of Fig. 5F, by developing and solving the ILP formulation in (3). In other words, the goal is to find a subnetwork of the initial curated network that exhibits the optimal fit to the data. The expectation after solving the ILP for the network in Fig. 6A is that the network in Fig. 5D must be obtained as the optimal solution.

### 4.3 Incorporating Feedback Paths in the Network Learning from Data

In the learning phase, care should be taken while considering the feedback paths. Since the network may present different responses at different time instances - which is the case as seen in Fig. 5F - implementing the constraints in (3) is not trivial for the nodes receiving incoming feedback inputs. In fact, it is very challenging to mathematically formulate such nodes in one step because such nodes need to be initialized and then updated when the LE data is considered. To solve this problem, we propose to duplicate the EE network, as shown in Fig. 7, and then connect these two identical networks using the feedback edges. Furthermore, we treat the intermediate and output nodes in the duplicate network as new nodes, as shown in Fig. 7 using the superscript ^*L*^ that refers to the late event. For instance, *x*_4_ and 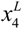 represent EE and LE activity variables, respectively, where 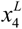 receives the feedback input initiated from *x*_6_. Note that if a node in the original network does not receive any feedback input, then its LE variable is equal to its EE variable, e.g., 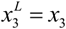. Moreover, the edges in the two identical network copies are labeled by the same decision variable *y_j_*, *j* ∈ *I_i_*, *i* = 1,…, *n_R_*, so that if *y_j_* = 0, then both edges are removed from the two network copies. For instance, the edges *x*_1_ —| *x*_4_ and 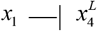 are both labeled by *y*_1_ in Fig. 7, so that *y*_1_ = 0 means that both edges are removed.

**Fig. 7.**
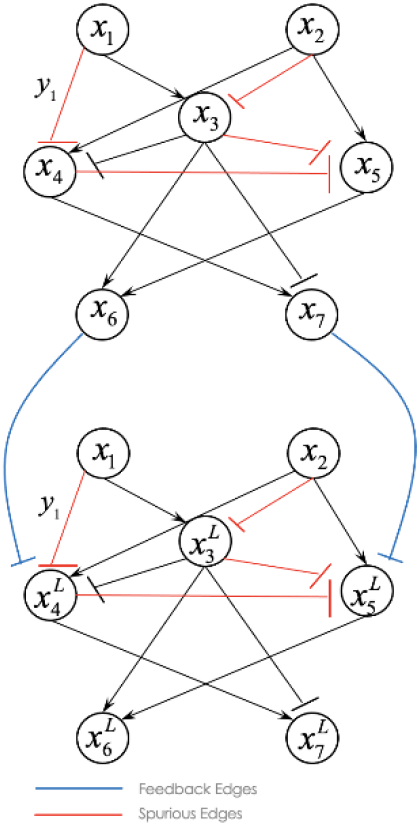
The duplicated network (bottom) of the original network (top), to incorporate the feedbacks during the network learning via ILP.

The ILP formulation for Fig. 7 is implemented using OPL (Optimization Programming Language), a high-level programming language, and is solved using the IBM ILOG CPLEX optimization studio [30], a commercial software that solves optimization problems. CPLEX found twelve optimal solutions, i.e., twelve *y* vectors, such that the numerical values of their learning objective function - computed using (2) - are equal to 0, which means 100% fitness. One of these optimal solutions is *y* =[0 1 1 0 1 1 0 1 1 0 1 1 1 1], which represents the network in Fig. 5D. This demonstrates the ability of the proposed approach in finding a subnetwork with the best fit to the data, while preserving the rules of the model of interest. How to handle other solutions is discussed in the next subsection.

One can similarly implement and solve the ILP formulation in (4) for network learning using the Model II given in Section 2.2, which is omitted here to avoid repetition.

### 4.4 Variations and Modifications

The proposed learning approach via ILP formulation and the introduced strategy to handle feedbacks are applicable to very large networks and capable of finding the exact optimal subnetworks with the best fitness percentage to the data. However, solutions of the ILP formulations may not be unique and multiple solutions may be obtained. On the other hand, we have observed that the results are usually highly correlated, indicating that the solutions are very similar. Therefore, one can examine the solutions and choose the one that meets a specific criterion, for example, being the closest to the published information in the literature.

It is also foreseeable that the resulting subnetwork solutions may be missing several interactions that existed in the initial network. To control the number of removed edges, one can add a penalty term to the learning objective function, for example, the one in (2), as follows:

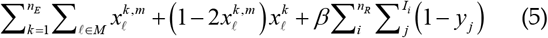

where *β* > 0 is a tunable penalty parameter that penalizes the objective function for each removed edge. Higher values of *β* may result in worse fitness to the data, while keeping more edges in the final learned subnetwork. Therefore, there can be a tradeoff between the number of removed edges and the data fitness percentage that we should keep in mind.

Finally, some of the interactions that are removed to obtain the optimal solution, may be well-known interactions that are experimentally confirmed by several groups of researchers. To prevent this from happening, one can add some additional constraints to force well-known edges of interest to remain in the network.

## 5 Conclusion

Transforming molecular networks into mathematically analyzable yet experimentally verifiable models is a major challenge in systems biology. The network models usually do not agree with the experimental measurements initially, specifically for literature-curated networks. The disagreement between model predictions and the experimental data can be due to the incompleteness of information resources, databases, and the literature used to construct the networks. Developing tools to learn the network models from empirical data is of high importance, since it improves the reliability of the models, and consequently increases the likelihood of confirming computational predictions in laboratory experiments. In this paper, we have presented two network models (Section 2) and have shown how the networks can be learned and calibrated using experimental data and via integer linear programming (Section 3), by minimizing the number of mismatches between the model predictions and the experimental data (Section 4).

Due to the feedback paths, modeling and analysis of molecular networks become more challenging. Because of the signal propagation delays introduced by the feedback mechanisms, network responses may change over time. Thus, the complex compensatory and regulatory mechanisms of feedbacks should be considered, while learning network models from data. Here, we have presented an efficient method for incorporating the effects of feedback paths in the learning algorithm (Fig. 7). As tested on an exemplary network (Fig. 6 and Fig. 7) the ILP formulation can effectively find the correct subnetwork from an initial network that has some spurious interactions. Different aspects of the proposed algorithm and relevant modifications are also discussed (Section 4.4).

Overall, the proposed network learning approach has promising potentials to reduce the gap between the literature-curated networks and the experimental data of the network, while incorporating complex interactions and biological mechanisms within the network such as feedbacks. This study is particularly important if computational analysis is going to be performed on the molecular network models associated with complex disorders, when some unknown molecular mechanisms are contributing to the development of the pathology. By providing more reliable networks with more accurate data prediction capabilities, the proposed approach of fitting the disease-associated molecular networks to the experimental data can assist in building better systems biology models for understanding the pathology, and eventually finding the best molecules in the network to target with novel therapeutic.

